# Components of TOR and MAP kinase signaling control chemotropism and pathogenicity in the fungal pathogen *Verticillium dahliae*

**DOI:** 10.1101/2022.06.20.496898

**Authors:** Vasileios Vangalis, Emmanouil A. Markakis, Michael Knop, Antonio Di Pietro, Milton A. Typas, Ioannis A. Papaioannou

**Affiliations:** Department of Genetics & Biotechnology, Faculty of Biology, National and Kapodistrian University of Athens, 15784, Athens, Greece; Laboratory of Mycology, Department of Viticulture, Vegetable Crops, Floriculture and Plant Protection, Institute of Olive Tree, Subtropical Crops and Viticulture, N.AG.RE.F., Hellenic Agricultural Organization - DIMITRA, 71307, Heraklion, Crete, Greece; Center for Molecular Biology of Heidelberg University (ZMBH), 69120, Heidelberg, Germany; German Cancer Research Center (DKFZ), DKFZ-ZMBH Alliance, 69120, Heidelberg, Germany; Departamento de Genética, Campus de Excelencia Internacional Agroalimentario ceiA3, Universidad de Córdoba, 14014, Córdoba, Spain

**Author notes:** Corresponding authors. (Milton A. Typas); (Ioannis A. Papaioannou). Email addresses (Vasileios Vangalis); (Emmanouil A. Markakis); (Michael Knop); (Antonio Di Pietro).

**Keywords:** autophagy, chemotropism, environmental sensing, mTOR signaling pathway, pathogenicity, *Verticillium dahliae*

## Abstract

Filamentous fungi can sense useful resources and hazards in their environment and direct growth of their hyphae accordingly. Chemotropism ensures access to nutrients, contact with other individuals (e.g., for mating), and interaction with hosts in the case of pathogens. Previous studies have revealed a complex chemotropic sensing landscape during host-pathogen interactions, but the underlying molecular machinery remains poorly characterized. Here we studied mechanisms controlling directed hyphal growth of the important plant-pathogenic fungus *Verticillium dahliae* towards different chemoattractants. We found that the homologs of the Rag GTPase Gtr1 and the GTPase-activating protein Tsc2, an activator and a repressor of the TOR kinase respectively, play important roles in hyphal chemotropism towards nutrients, plant-derived signals, and heterologous α-pheromone of *Fusarium oxysporum*. Furthermore, important roles of these regulators were identified in fungal development and pathogenicity. We also found that the mitogen-activated protein kinase (MAPK) Fus3 is required for chemotropism towards nutrients, while the G protein-coupled receptor (GPCR) Ste2 and the MAPK Slt2 control chemosensing of plant-derived signals and α-pheromone. Our study establishes *V. dahliae* as a suitable model for the analysis of fungal chemotropism and discovers new components of chemotropic signaling, during growth and host-pathogen interactions of *V. dahliae*.

## 1. Introduction

Filamentous fungi grow and colonize their substrates by polarized tip extension and lateral branching of their hyphae (Riquelme, 2013). Since the necessary resources for their survival are heterogeneously distributed in their habitats, fungi have evolved mechanisms for sensing the environment and responding by directed growth (Brand and Gow, 2009; Leeder et al., 2011). This results from dynamic re-orientation of hyphal growth towards or away from stimuli such as nutrients, toxic substances, other individuals (e.g., during mating and vegetative fusion), and host organisms (Lombardi et al., 2018; Moreno-Ruiz et al., 2020; Sridhar et al., 2020; Turrà et al., 2015; Vangalis et al., 2021a). In the case of fungal pathogens, chemotropic sensing and growth play prominent roles during host-pathogen interactions (Turrà et al., 2016). However, the underlying mechanisms of sensing and processing chemotropic signals are still not fully understood (Turrà et al., 2016; Van Dijck et al., 2017).

Various classes of membrane proteins such as non-transporting receptors, transceptors, and G protein-coupled receptors (GPCRs) have important roles in nutrient sensing (Van Dijck et al., 2017). Studies in plant-pathogenic *Fusarium* species have implicated the GPCR Ste2, a homolog of the alpha-factor pheromone receptor of *Saccharomyces cerevisiae*, in chemotropic sensing of the host plant (Nordzieke et al., 2019; Sridhar et al., 2020; Turrà et al., 2015). The signal is then transduced to the mitogen-activated protein kinase (MAPK) Mpk1 (Slt2 in yeast), a component of the cell wall integrity (CWI) signaling pathway, to mediate the chemotropic response (Nordzieke et al., 2019; Sridhar et al., 2020; Turrà et al., 2015).

The CWI MAPK cascade is required for remodeling of the fungal cell wall and ensures its integrity under cell wall stress. Deletion of Slt2 homologs of filamentous fungi results in defects in hyphal growth, hypersensitivity to cell wall-damaging agents, and often reduced pathogenicity against plant or animal hosts (reviewed by Zhao et al. 2007; Turrà et al. 2014; Jiang et al. 2018). Apart from its role in the chemotropic response to plant signals, Mpk1 in *F. oxysporum* has also been implicated in the response to α-pheromone as a cell density-dependent autocrine signal to control germination of conidia (Turrà et al., 2015; Vitale et al., 2019). Furthermore, the homolog of *Neurospora crassa* has been shown to participate, together with the Fus3 homolog, in an oscillatory mechanism that mediates communication of anastomosis tubes during vegetative hyphal fusion (Fischer and Glass, 2019; Fleissner et al., 2009). Moreover, in *F. oxysporum,* the MAPK homolog of Fus3 is required for the chemotropic response to different nutrients in a Ste2-independent manner (Turrà et al., 2015). In addition to these functions, Fus3 homologs play important roles in fungal development and pathogenicity (reviewed by Zhao et al. 2007; Turrà et al. 2014; Jiang et al. 2018).

In eukaryotes, Target Of Rapamycin (TOR) kinase complexes respond to multiple environmental cues (Dobrenel et al., 2016; Workman et al., 2014) to regulate cellular development, proliferation, and survival by controlling metabolic processes, nutrient uptake, autophagy, as well as protein production and degradation (Chantranupong et al., 2015; Saxton and Sabatini, 2017; Workman et al., 2014). In filamentous fungi, the TOR signaling pathway was shown to be involved in the regulation of nutrient sensing, hyphal growth and morphogenesis, cell cycle regulation, and stress response (Fitzgibbon et al., 2005; Li et al., 2019; Weisman, 2016; Yu et al., 2014). The GTPases Rag1/2 and Rheb, as well as their respective repressors GATOR1 and TSC1/2, are important regulators of the TOR kinase complex 1 (TORC1) in higher eukaryotes, and they are conserved in fungi (i.e., GTPases Gtr1/2 and Rhb1, and GAP repressors SEACIT and TSC1/2, respectively) (Wolfson and Sabatini, 2017). Characterization of the Gtr1/2 complex in *S. cerevisiae* revealed its participation in the activation machinery of TORC1 in response to amino acids or other nitrogen sources (Binda et al., 2009; Kira et al., 2016, 2014), and this was further supported by evidence from *Schizosaccharomyces pombe* (Valbuena et al., 2012). On the other hand, deletion of *tsc1* or *tsc2* in *S. pombe* led to hyperactivation of TORC1 and a reduction of amino acid uptake (Matsumoto et al., 2002; Slegtenhorst et al., 2004).

The ascomycete *Verticillium dahliae* is a soilborne plant pathogen that causes economically devastating wilt disease on a wide range of commercially important plants and crops (Pegg, G.F and Brady, B.L., 2002). The infection cycle of *V. dahliae* starts with germination of microsclerotia (i.e., resting structures) induced by plant root exudates, penetration of roots by elongating hyphae, and colonization of the host’s vascular system (Klosterman et al., 2009). The mechanisms used for sensing and growth towards plant-derived signals are currently unknown.

In this study, we used *V. dahliae* as a fungal model to elucidate the molecular machinery for environmental sensing and chemotropism towards nutrients and plant hosts. For this, we examined two members of the TOR signaling pathway, Gtr1 and Tsc2, that have not been studied previously in filamentous fungi, as well as the MAPKs Fus3 and Slt2, and the GPCR Ste2. Our study provides new insights into the fungal nutrient and host sensing machinery, establishes *V. dahliae* as a suitable model for such analyses, and further elucidates important pleiotropic roles of MAPK and TOR signaling in fungal plant pathogens.

## 2. Materials and Methods

### 2.1. Strains and culture conditions

All *V. dahliae* strains and plasmids that were used and constructed in this study are listed in Supplementary Tables 1 and 2, respectively (Lu et al., 1994; Nguyen et al., 2008; Paz et al., 2011). Culture media and conditions, preparation, handling, and maintenance of monoconidial strains have been described previously (Papaioannou et al., 2013; Vangalis et al., 2021a).

### 2.2. Gene deletion and complementation of knockout strains

Deletion strains Δ*gtr1* and Δ*tsc2*, as well as double deletion strains Δ*fus3* Δ*slt2* and Δ*fus3* Δ*ste2* were constructed in this study in the background of the wild-type *V. dahliae* isolate 123V (Supplementary Table 1). For this, we used our optimized *Agrobacterium tumefaciens*-mediated transformation protocol and constructed plasmids for double homologous recombination, as previously described (Vangalis et al., 2020). All knockout mutants were validated by PCR using gene-specific primers and Southern hybridization analyses (Supplementary Fig. 1). For complementation of the knockout strains, the corresponding wild-type genes with their flanking regions were re-introduced, according to our previously reported procedures (Vangalis et al., 2020). We also included in parts of our study *V. dahliae* 123V deletion mutants Δ*fus3*, Δ*slt2*, and Δ*ste2*, which were previously reported (Vangalis et al., 2021a).

### 2.3. Quantitative analysis of chemotropism

Chemotropic responses were quantified using a plate assay previously employed in *Fusarium oxysporum* (Turrà et al., 2015) with minor modifications. Freshly obtained *V. dahliae* conidia were embedded in 5 mL of three times diluted Czapek-Dox Minimal Medium (MM/3) supplemented with 0.5% agar (w/v), at a final concentration of 2.5×10^6^ per mL, which was poured into a Petri dish. A central scoring line was drawn on the bottom of the plate, and two parallel wells were cut into the medium on both sides, at 6 mm distance from the scoring line. Then, 50 μL of the test compound solution and of the solvent control were added to the wells at the two sides of the scoring line, respectively. Tested compounds and standard concentrations used were: sodium aspartate (Asp) and methionine (Met) at 295 mM; ammonium nitrate (NH_4_^+^), glucose (Gluc), glycerol (Glyc), and sodium glutamate (Glu) at 100 mM; and pectin (Pec) at 1% (w/v) (all from Sigma Aldrich, Saint Louis, MO, USA). Sterile water or methanol were used as solvent controls, as appropriate. To measure the chemotropic response towards host plants, the root of a 2-week-old tomato seedling was placed directly on one of the wells. Lyophilized peptides of synthetic *F. oxysporum* α-pheromone (Turrà et al., 2015) were dissolved in 50% (v/v) methanol in water and assayed at a concentration of 378 μM. To test the effect of different treatments on the chemoattractant activity of *F. oxysporum* α-pheromone or tomato root exudates, the peptide and the exudates were incubated for 10 min at 100 °C or treated with 1 mg/mL proteinase K (Sigma Aldrich, Saint Louis, MO, USA). Plates were incubated at 25 °C in the dark for 14 h, and chemotropism of conidial germ tubes was then recorded as described (Turrà et al., 2015) using an Olympus (Olympus Corporation, Shinjuku City, Tokyo, Japan) binocular microscope (200× magnification). The chemotropic index used to quantify chemotropism (Fig. 1A) was calculated by the formula [(germ tubes pointing the stimulus – germ tubes pointing the solvent)/(total number of germ tubes) × 100] (Turrà et al., 2015). A total of 400 hyphal tips were scored for each test compound. All experiments were performed at least twice. Statistical analysis was conducted using *t*-test.

**Fig. 1.**
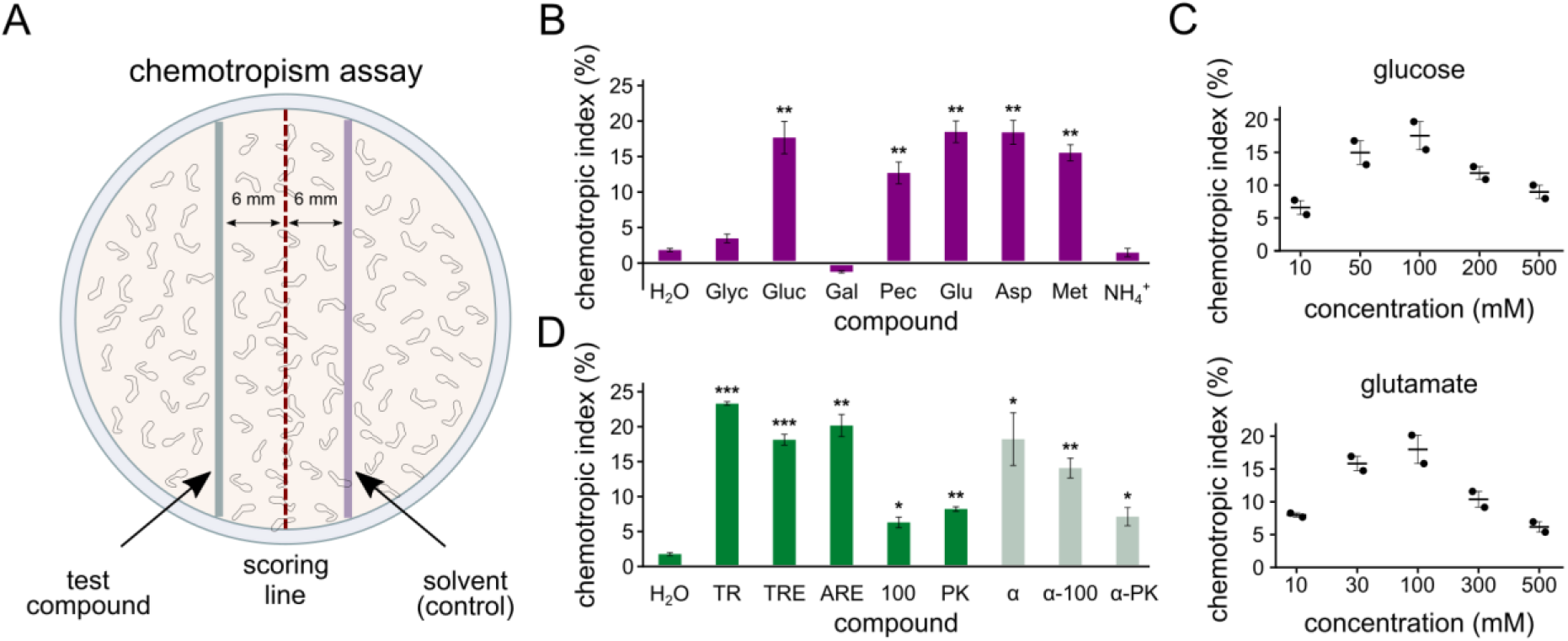
Optimization of a chemotropism assay in *V. dahliae*. **A.** Schematic representation of the chemotropism assay (modified from Turrà et al. 2015). **B.** Chemotropism of the wild-type strain 123V towards the nutrient sources glycerol (Glyc), glucose (Gluc), galactose (Gal), pectin (Pec), glutamate (Glu), aspartate (Asp), methionine (Met), and ammonium nitrate (NH_4_^+^). **C.** Dose-dependent chemotropic response to the indicated concentrations of glucose and glutamate. **D.** Chemotropism of the wild-type strain 123V towards tomato roots (TR), tomato root exudate (TRE), *Arabidopsis* root exudate (ARE), α-pheromone (α), or towards tomato root exudate or α-pheromone after boiling (100 and α-100, respectively) or treatment with proteinase K (PK and α-PK, respectively). Bars (in **B-D**): SD. Statistical significance of differences from the control condition (H_2_O) in **B** and **D** was tested with Student’s *t*-test (* *p*<0.05, ** *p*<0.01, *** *p*<0.001).

### 2.4. Western blot analyses

To study MAPK phosphorylation in response to different compounds, 5×10^6^ conidia per mL were allowed to germinate for 24 h in PDA medium at 25 °C and 170 rpm. Samples of fungal mycelium were collected before (time 0) and after the addition of each compound. Protein extraction was performed with a lysis buffer containing 0.2 M NaOH and 0.2% (v/v) β-mercaptoethanol, followed by precipitation with 7.5% trichloroacetic acid (TCA) (Méchin et al., 2007). Western blotting was performed as described (Segorbe et al., 2017). Protein samples were resolved in 10% SDS-polyacrylamide gels and transferred to nitrocellulose membranes using the Trans-Blot Turbo RTA Midi Nitrocellulose Transfer Kit (#1704271 from Bio-Rad Laboratories, Hercules, CA, USA). Phosphorylation of MAPKs Slt2 and Fus3 was detected using a rabbit antiPhospho-p44/42 MAPK (Erk1/2) antibody (Thr202/Tyr204; #4370 from Cell Signaling Technology, Danvers, MA, USA). A mouse anti-α-tubulin antibody (#T9026 from Sigma Aldrich, Saint Louis, MO, USA) was used for the loading controls. Visualization of the corresponding bands was performed using the ECL™ Select western blotting detection reagent (GE Healthcare, Chicago, IL, USA) in a LAS-3000 detection system (Fujifilm España, Barcelona, Spain).

### 2.5. Plant pathogenicity assays

Plant pathogenicity bioassays were performed according to previously published protocols (Markakis et al., 2014; Vangalis et al., 2020). Briefly, eggplant seedlings at the one true leaf stage were drenched with 20 mL of conidial suspensions (5.0×10^6^ conidia per mL) or sterile water (mock infection controls). Treated plants were incubated at 25 °C with a 12-h light-dark cycle. Assessment of disease severity at each time point (up to 31 days) and determination of relative AUDPC and plant growth parameters was performed as previously described (Markakis et al., 2014; Vangalis et al., 2020). Fungal re-isolation was performed from sections along the stem of each treated plant (three chips from nine randomly selected plants per treatment) as previously described (Markakis et al., 2014). The re-isolation ratio was expressed as the percentage of xylem chips from each treatment that exhibited fungal growth.

### 2.6. Morphological characterization and stress response of V. dahliae strains

Morphological and physiological characterization of fungal strains, and assessment of their stress tolerance were performed as previously described (Vangalis et al., 2021c). All experiments were performed at least in triplicate. To determine the frequency of conidial germination, 100 conidia per strain and replicate were analyzed. Strains were exposed to a variety of oxidative agents (H_2_O_2_, paraquat, iprodione, farnesol), substances that induce osmotic stress (NaCl, sorbitol), cell wall damaging factors (amphotericin B obtained from Biosera, Nuaille, France; fluconazole from Pfizer, Brooklyn, NY, USA; calcofluor white M2R, Congo red), an inhibitor of TOR kinase (sirolimus from Cayman Chemical, Ann Arbor, MI, USA), and fungicides (azoxystrobin, cymoxanil). All were purchased from Sigma Aldrich (Saint Louis, MO, USA) unless otherwise indicated. For quantification of sensitivity to the tested compounds, relative growth inhibition was calculated by the formula [(colony diameter on CM − colony diameter in stress condition)/(colony diameter on CM) × 100)].

## 3. Results

### 3.1. A quantitative assay for the analysis of chemotropism in V. dahliae

For the purposes of our study, we first optimized a quantitative chemosensing assay for *V. dahliae* on agar plates, based on a procedure that was previously used in *F. oxysporum* (Turrà et al., 2015). Our optimized assay (described schematically in Fig. 1A; details in paragraph 2.3) exposes *V. dahliae* germlings to gradients of diffusing substances and enables microscopic recording of the growth orientation of hyphal tips. When germlings of the wild-type strain 123V were tested against a variety of carbon and nitrogen sources, we found glucose, pectin, and the amino acids glutamate, aspartate, and methionine to induce strong positive chemotropic responses, in contrast to glycerol, galactose, and ammonium nitrate (Fig. 1B). Our further analyses of responses to glucose and glutamate revealed that hyphal chemotropism is dose-dependent, with particularly low and high concentrations both failing to induce strong reactions (Fig. 1C). Hyphal tips were significantly attracted by tomato roots, as well as by root exudates of tomato and *Arabidopsis* plants that were tested (Fig. 1D). This directed hyphal growth was largely reduced when exudates were boiled or treated with proteinase K. Although *V. dahliae* is widely regarded as an asexual fungus, it exhibited significant chemotropic responses to synthetic α-pheromone of *F. oxysporum*, which were also reduced upon treatment with proteinase K (Fig. 1D).

### 3.2. Differential roles of the MAPKs Fus3 and Slt2 in chemotropism of V. dahliae towards nutrients, α-pheromone, and plant signals

The involvement of MAPK signaling in environmental sensing in other organisms (Turrà et al., 2014, 2016) motivated us to analyze further the *V. dahliae* homologs of MAPKs Fus3 and Slt2. Deletion of *fus3* led to markedly reduced production of microsclerotia (Fig. 2A), reduced growth, compromised sporulation (Fig. 2B), and increased sensitivity to oxidative stress (Supplementary Fig. 2). The Δ*slt2* mutant also exhibited complete inability to produce microsclerotia, drastic reduction of aerial mycelium (Fig. 2A), defective sporulation (Fig. 2B), and significant growth inhibition by chemicals that induce cell wall stress (Supplementary Fig. 2). Double deletion of *fus3* and *slt2* caused the combined defects of the single mutants as well as an additional defect in conidial germination (Fig. 2A-B).

**Fig. 2.**
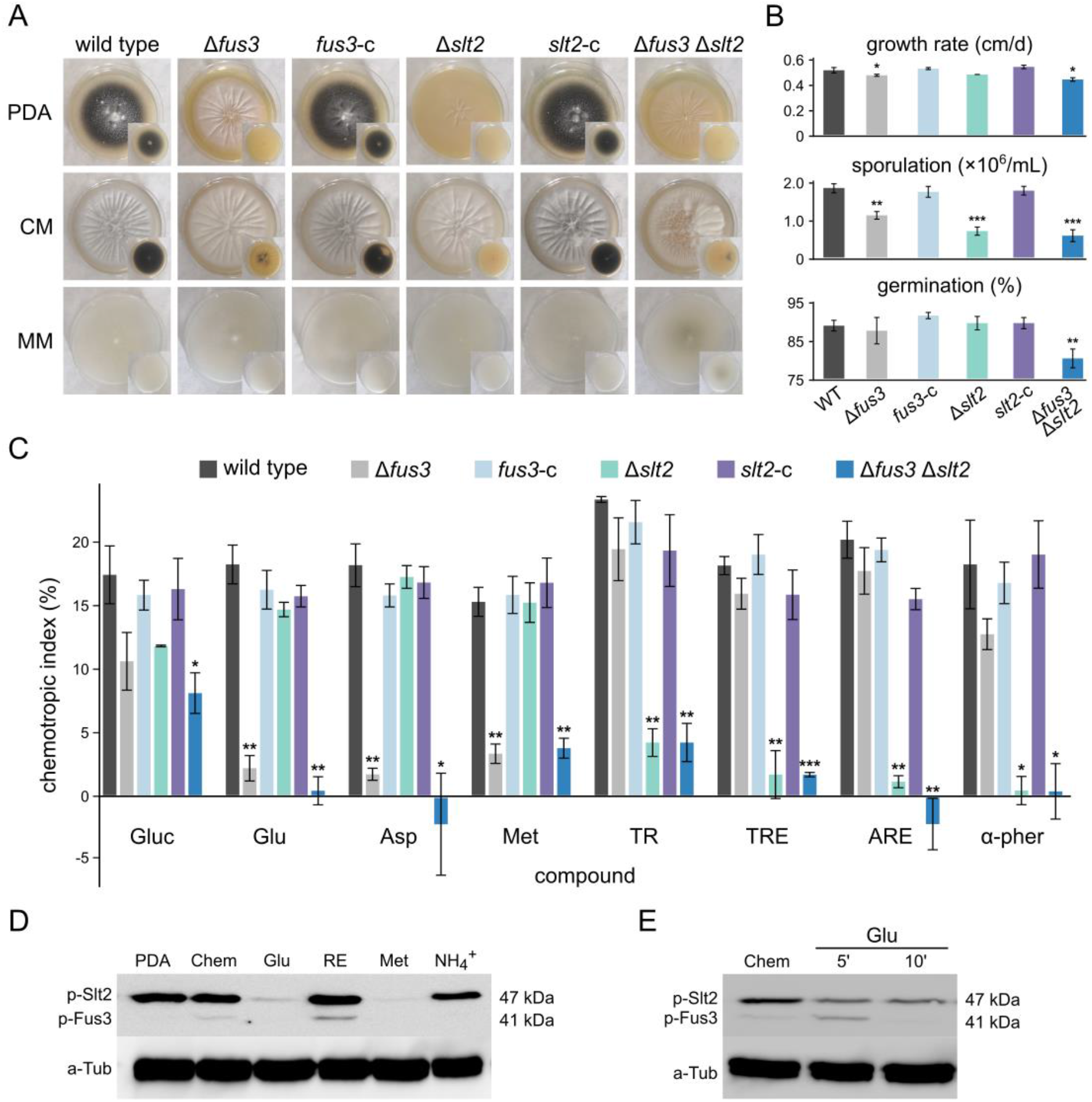
The MAP kinases Fus3 and Slt2 are involved in *V. dahliae* chemotropism towards environmental nutrients and hosts/α-pheromone, respectively. **A.** Morphological features of *V. dahliae* 123V deletion mutants of *fus3* and *slt2* 35 days after inoculation on the indicated agar media. *Fus3*-c and *slt2*-c are the corresponding complemented strains. **B.** Growth rate, sporulation, and conidial germination frequency of the tested strains. **C.** Chemotropic responses of the studied mutants towards various environmental stimuli (abbreviations as in Fig. 1). In **B-C**, bars represent the SD of replicates, and statistical significance of differences from the wild type was tested with Student’s *t*-test (\**p*<0.05, \*\**p*<0.01, \*\*\**p*<0.001). **D.** Western blot analysis of the phosphorylation levels of Slt2 after 15 minutes of exposure to the indicated stimuli (abbreviations as in Fig. 1; PDA: potato dextrose broth; Chem: chemotropic assay medium – time 0’ control). **E.** Western blot analysis of the phosphorylation levels of Slt2 at the indicated time points after addition of glutamate to the culture medium.

We used the chemotropism assay to test these deletion mutants for the ability to sense and orient their hyphal tips towards nutrients or hosts in their environment. The MAPK Fus3 was necessary for normal chemotropism towards nutrients, while Slt2 was required for chemotropism towards root exudates and synthetic *F. oxysporum* α-pheromone peptide (Fig. 2C). In line with this, the double mutant Δ*fus3* Δ*slt2* failed to respond to any of the tested stimuli (Fig. 2C). Wild-type behavior was fully rescued by complementation of our deletion mutants with the respective wild-type alleles (Fig. 2C).

We next determined the MAPK phosphorylation levels in germlings exposed for 15 min to various stimuli. Immunoblot analysis revealed significant dephosphorylation of Slt2 upon exposure to the nutrient chemoattractants glucose and methionine, but not to root exudates (Fig. 2D). Interestingly, exposure to ammonium nitrate, a nutrient source that does not induce a chemotropic response did not alter the phosphorylation levels of Slt2 (Fig. 2D). A time-course experiment revealed that Slt2 dephosphorylation in response to glucose was already detectable 5 min upon exposure (Fig. 2E).

### 3.3. Fus3, but not Slt2, is required for pathogenicity of V. dahliae

Our finding that Slt2 is required for chemotropism of *V. dahliae* towards the host plant led us to investigate the hypothesis that this MAPK might be important for pathogenicity, as previously shown for Fus3 in a different *V. dahliae* isolate (Rauyaree et al., 2005). We therefore tested the ability of the deletion mutants to cause disease to eggplant, a particularly susceptible host of *V. dahliae* that allows convenient and reproducible scoring of disease symptoms. Our results confirmed that deletion of *fus3* severely compromises pathogenicity, since plants treated with the single Δ*fus3* and the double Δ*fus3* Δ*slt2* deletion mutant showed only very mild symptoms (Fig. 3A-D). In contrast, virulence of Δ*slt2* was indistinguishable from that of the wild-type strain (Fig. 3A-D). At the end of the experiment (31 days after inoculation), measurement of the plant fresh weight and fungal re-isolation from xylem chips yielded consistent results, with Δ*fus3* and Δ*fus3* Δ*slt2* failing to reduce plant weight and rarely being recovered from the xylem of inoculated plants, whereas the results of the Δ*slt2* mutant were very similar to those of the wild-type strain (Fig. 3D).

**Fig. 3.**
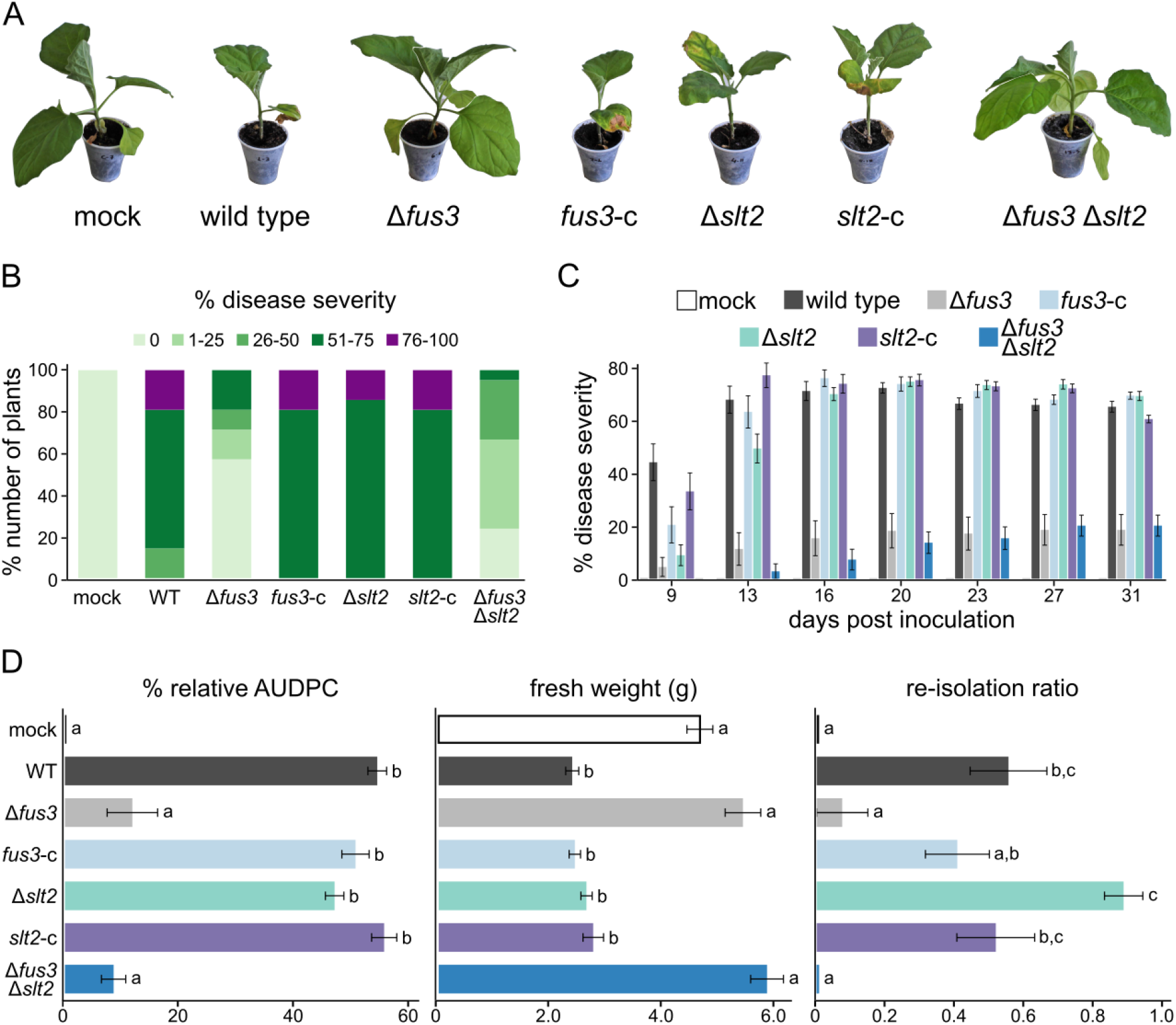
Fus3, but not Slt2, is required for pathogenicity of *V. dahliae* on eggplant. **A.** Representative examples of plants inoculated with the wild-type strain (123V), as well as the indicated deletion mutants and complemented (*fus3*-c and *slt2*-c) strains were imaged 31 days post-inoculation (d.p.i.). **B.** Average disease severity at 31 d.p.i. (21 eggplant seedlings/strain). Non-infected plants (mock) served as controls. Percentages in different colours represent different levels of disease severity**. C.** Time-course analysis of disease severity. **D.** Mean relative area under disease progress curve (AUDPC) score for each strain (left), average plant fresh weight (center), and fungal re-isolation ratio (right) at the end of the infection experiment (31 d.p.i.). In **C-D**, error bars represent the SE of replicates, and statistical significance of differences between strains was tested by one-way ANOVA followed by Tukey’s post hoc tests. Bars marked with the same letter do not differ significantly (*p*≤0.05).

### 3.4. Ste2 is required for chemotropism towards α-pheromone and the host plants

Binding of mating peptide pheromone to its cognate receptor (i.e., Ste2 and Ste3 in yeast) is essential for sensing mating partners and constitutes the first step in the mating pathway. Previously, the Ste2 homolog of the asexual fungal pathogen *F. oxysporum* was shown to be required for chemotropism towards α peptide pheromone and tomato root exudates (Turrà et al., 2015). We therefore tested whether Ste2 may also be involved in chemotropic sensing of *V. dahliae*.

Unlike Δ*fus3*, the Δ*ste2* mutant exhibited no detectable defects in morphology, physiology, stress response, or pathogenicity, while the Δ*fus3* Δ*ste2* double deletion mutant displayed a similar phenotype as Δ*fus3* (Fig. 4A-C, Supplementary Fig. 2). Chemotropism assays with these mutants demonstrated that Ste2 is not required for chemosensing of different nutrients (Fig. 4D), a process that is dependent on Fus3. However, Ste2 was required for chemotropism towards tomato root exudates or *F. oxysporum* α-pheromone (Fig. 4D), similar to Slt2. The Δ*fus3* Δs*te2* double deletion mutant showed a severely reduced chemoresponse to all tested compounds (Fig. 4D), similar to the Δ*fus3* Δ*slt2* strain. Western blot analysis revealed differences in Slt2 phosphorylation dynamics between the wild-type and the Δs*te2* mutant upon exposure to chemotropic stimuli (Fig. 4E). Collectively, these results suggest a possible link between Ste2 and the Slt2 MAPK cascade in fungal chemosensing and chemotropism.

**Fig. 4.**
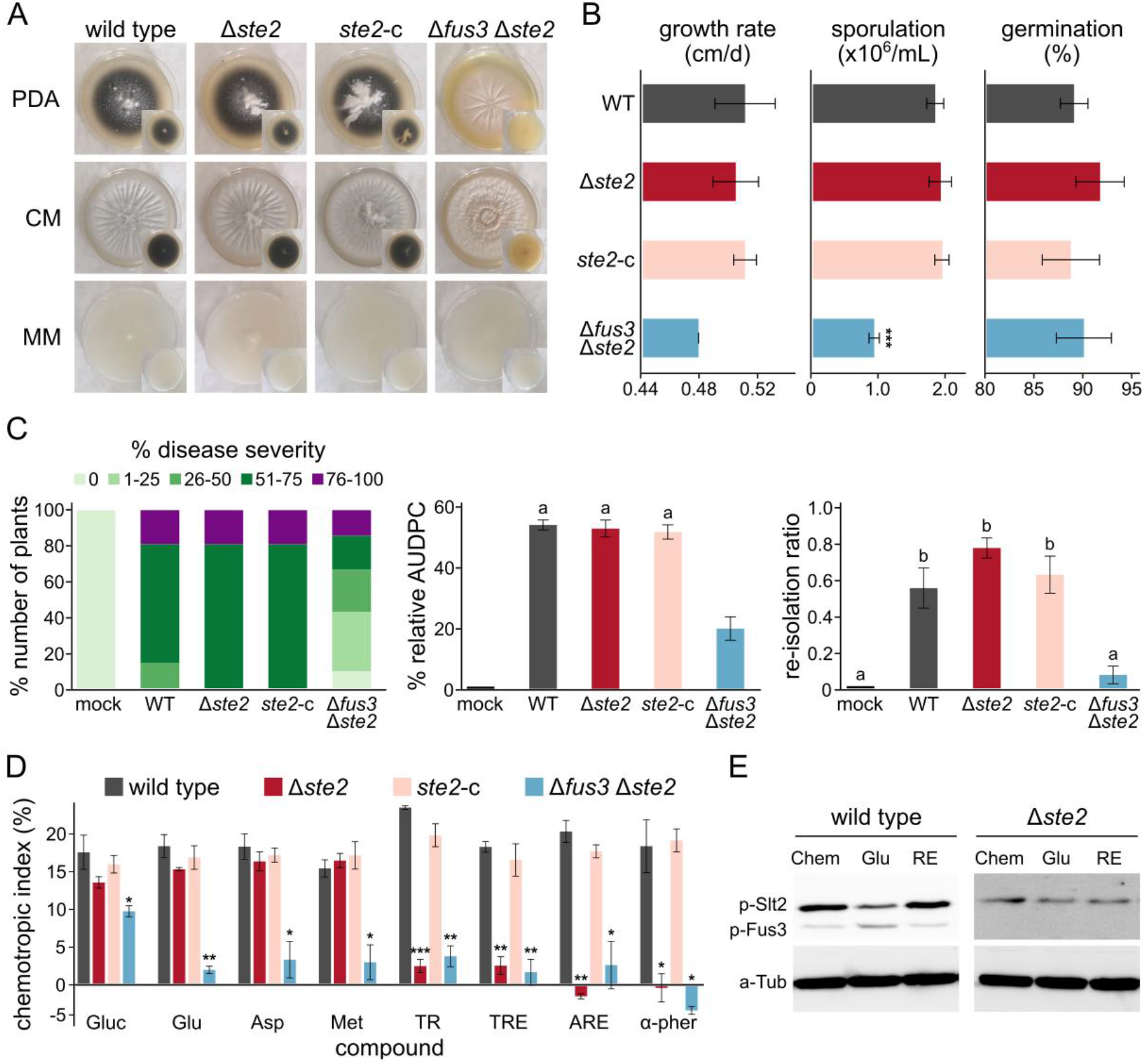
Ste2 is involved in chemotropism towards hosts and α-pheromone. **A.** Morphological characteristics of *V. dahliae* 123V deletion mutants Δ*ste2* and Δ*fus3* Δ*ste2* (*ste2*-c: complemented strain) 35 days after inoculation on the indicated agar media. **B.** Growth rate, sporulation, and conidial germination frequency of the same strains. **C.** Average disease severity (left), mean relative area under disease progress curve (AUDPC) score (middle), and fungal re-isolation ratio (right) for the same *V. dahliae* strains 31 days after inoculation (21 eggplant seedlings/strain). Error bars: SE. Statistical significance of differences between strains was tested by one-way ANOVA followed by Tukey’s post hoc tests. Bars marked with the same letter did not differ significantly (*p*≤0.05). **D.** Chemotropic responses of the studied mutants towards various environmental stimuli (abbreviations as in Fig. 1). In **B** and **D**, bars indicate the SD of replicates, and statistical significance of differences from the wild type was tested with Student’s *t*-test (* *p*≤0.05, ** *p*≤0.01, *** *p*≤0.001). **E.** Western blot analysis of the phosphorylation levels of Slt2 after addition of 100 mM glutamate (Glu; 5 min) or tomato root exudate (RE; 10 min); Chem: chemotropic assay medium – time 0’ control.

### 3.5. The TOR signaling regulators Gtr1 and Tsc2 mediate chemotropism towards nutrients and plant signals

The TOR kinase in eukaryotes is a central molecular hub where various environmental signals converge (González and Hall, 2017). Since deletion of *tor* is lethal in *V. dahliae* (Fang et al., 2017), we analyzed deletion mutants of two conserved regulators of the TOR pathway, the positively acting RAG GTPase Gtr1/2 and the GTPase activating protein (GAP) Tsc1/2, which acts as a repressor of the TOR pathway (Gao et al., 2002; Valbuena et al., 2012). By using BLAST searches with the amino acid sequences of *S. cerevisiae* Gtr1 and *S. pombe* Tsc2 as queries, we identified in the *V. dahliae* genome (reference strain Ls.17) single homologs of *gtr1* (VDAG_09326, 375 aa; 80% query coverage, 46% identity, 63% similarity) and *tsc2* (VDAG_09981, 1617 aa; 44% query coverage, 39% identity, 58% similarity). We then generated Δ*gtr1* and Δ*tsc2* knockout mutants (Supplementary Fig. 1), both of which displayed reduced production of microsclerotia and limited formation of aerial mycelium (Fig. 5A), as well as lower growth rates and reduced sporulation on CM in comparison to the wild-type strain (Fig. 5B). Moreover, the Δ*tsc2* mutant exhibited defects in conidial germination (Fig. 5B). Both deletion strains showed increased sensitivity to hyperosmotic stress and to the antifungal amphotericin B, Δ*gtr1* had impaired growth in the presence of fluconazole, while Δ*tsc2* was significantly inhibited by farnesol and paraquat (Supplementary Fig. 2).

**Fig. 5.**
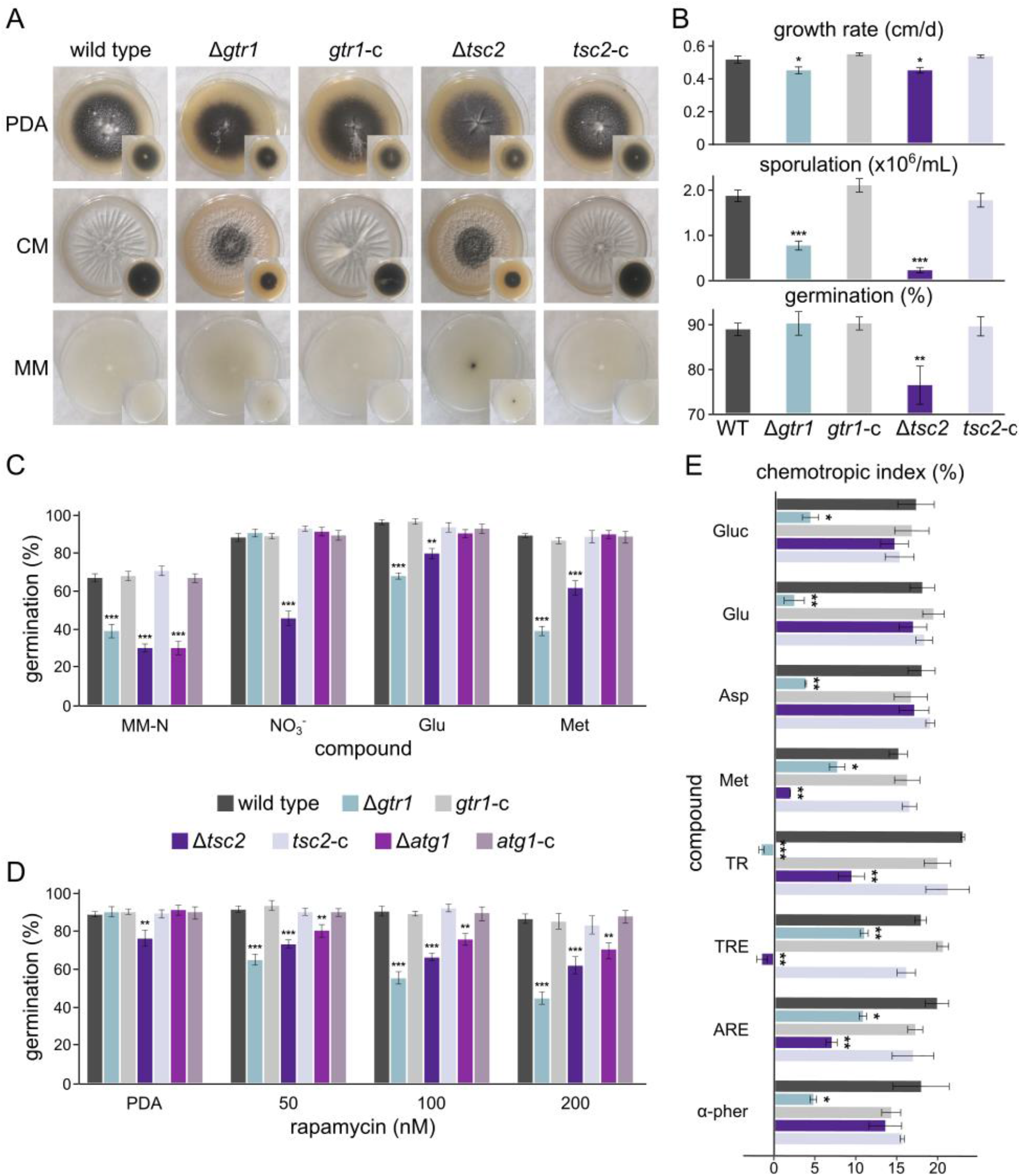
Chemotropism is mediated by regulators of the TOR signaling pathway. **A.** Morphology of the *V. dahliae* wild-type strain (123V), Δ*gtr1*, Δ*tsc2*, and their complemented (*gtr1*-c and *tsc2*-c) strains after 35 days of growth on the indicated agar media. **B.** Growth rate, sporulation, and conidial germination frequency of the same strains. **C.** Conidial germination frequency of the indicated strains after 20 h of nitrogen starvation (MM-N) or incubation in MM supplemented with sodium nitrate (NO_3_^-^), glutamate (Glu), or methionine (Met). **D.** Conidial germination frequency of the indicated strains after 20 h of incubation in PDB supplemented with rapamycin. **E.** Chemotropic responses of the indicated strains 14 h after initial exposure to the indicated stimuli (for abbreviations see Fig. 1). In **B-E,** bars indicate the SD of the replicates. Statistical significance of the differences from the wild-type strain was tested with Student’s *t*-test (* *p*≤0.05, ** *p*≤0.01, *** *p*≤0.001).

To confirm the role of these genes in TOR signaling of *V. dahliae*, we tested sensitivity of Δ*gtr1* and Δ*tsc2* to factors that induce autophagy and compared their phenotypes to that of the Δ*atg1* mutant which is impaired in autophagy (Vangalis et al., 2021b). The ability of conidia to germinate in the absence of nitrogen sources was significantly reduced in all deletion mutants compared to the wild-type strain (Fig. 5C), which is consistent with a defective autophagy in the knockout mutants. Addition of different nitrogen sources fully rescued germination in both the wild-type strain and Δ*atg1,* but only partially in Δ*gtr1* and Δ*tsc2* (Fig. 5C). Treatment of conidia with rapamycin, a known inhibitor of TORC1, caused a significant and concentration-dependent reduction in germination of Δ*gtr1*, Δ*tsc2*, and Δ*atg1*, but not in the wild-type and the complemented strains (Fig. 5D). These findings indicate that *V. dahliae* Gtr1 and Tsc2 are involved in regulation of TOR activity and induction of autophagy, depending on the levels of nutrient availability, consistent with their roles in other organisms (Wolfson and Sabatini, 2017).

We next asked whether Gtr1 and Tsc2 participate in the chemotropic response to nutrients and host signals. The Δ*gtr1* mutant showed a significantly reduced ability to re-direct hyphal growth towards all tested stimuli including nutrients, α-pheromone, and tomato root exudate, while the Δ*tsc2* mutant showed a reduced chemotropic response to root exudate and methionine (Fig. 5E). Western blot analysis detected altered phosphorylation patterns of Slt2 upon addition of chemoattractant compounds in the Δ*gtr1* and Δ*tsc2* mutants as compared to the wild-type strain (Supplementary Fig. 3), indicating a possible crosstalk between TOR and MAPK signaling in this process.

### 3.6. TOR signaling and autophagy contribute to pathogenicity in V. dahliae

Pathogenicity bioassays in eggplant seedlings were performed to investigate the possible involvement of Gtr1 and Tsc2 in *V. dahliae* virulence. A small but significant decrease in the frequency of severe disease symptoms was observed in plants infected with the Δ*gtr1* and Δ*tsc2* mutants in comparison to the wild-type strain (Fig. 6A-B). A time-course comparison of symptom development revealed a delay in emergence of symptoms in the deletion mutants, with differences from the wild-type strain being more pronounced in the first 16 days after inoculation (Fig. 6C). Calculation of the average AUDPC scores over the 31 days of the experiment revealed a reduction in the Δ*gtr1* and Δ*tsc2* mutants by 33.3% and 34.8%, respectively, compared to the wild type (Fig. 6D). Moreover, the average fresh weight of plants infected with Δ*gtr1* and Δ*tsc2* was significantly lower than that of the plants treated with the wild-type strain (Fig. 6D). When we investigated the presence of the fungus within the plant tissues, the re-isolation ratio of Δ*tsc2* was significantly lower than that of the wild-type strain (Fig. 6D).

**Fig. 6.**
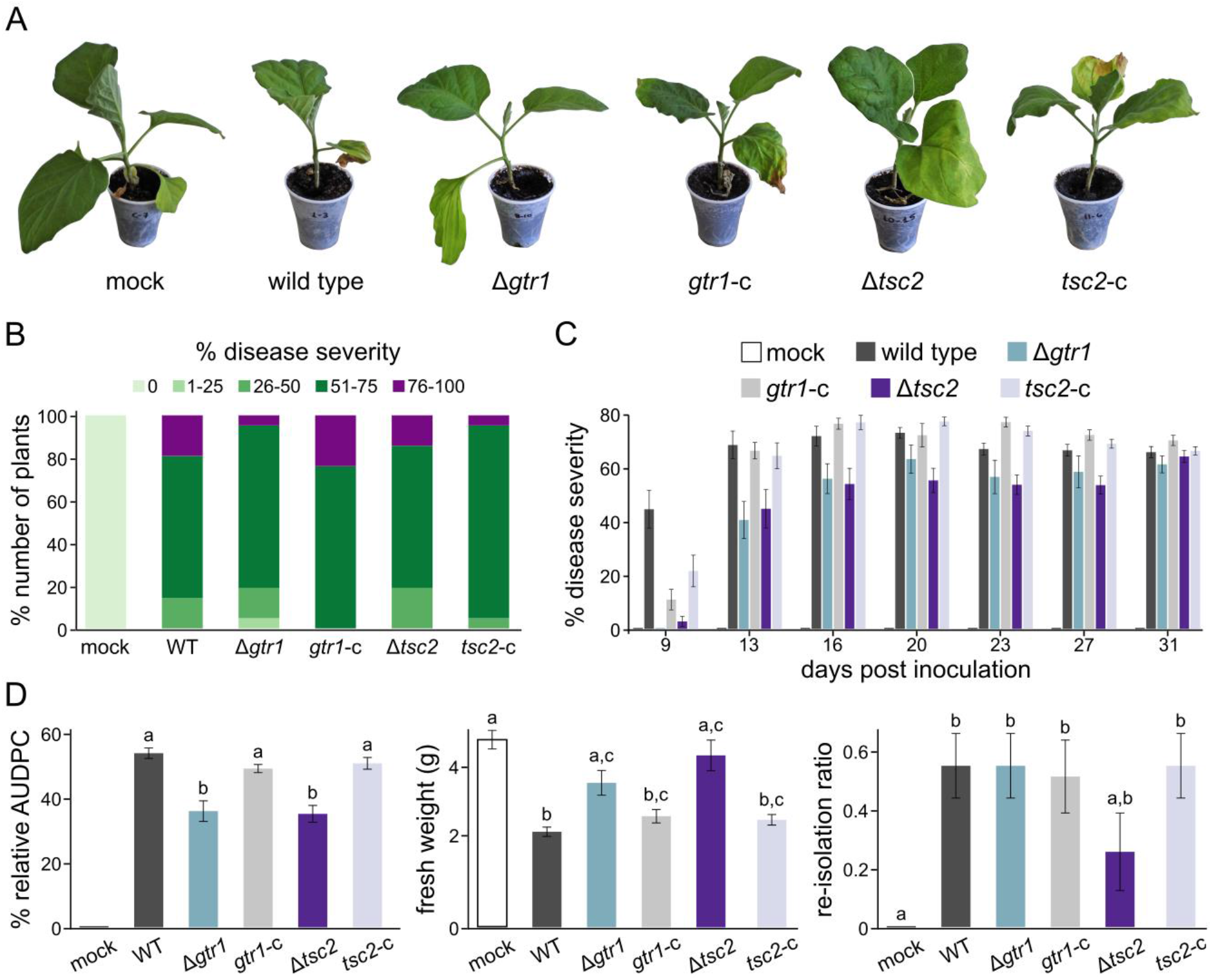
The TOR signaling regulators Gtr1 and Tsc2 contribute to pathogenicity of *V. dahliae*. **A.** Representative examples of eggplant plants inoculated with the wild-type strain (123V), the deletion mutants, or the complemented (*gtr1*-c and *tsc2*-c) strains at 31 days post-inoculation (d.p.i.). **B.** Average disease severity at 31 d.p.i. (21 eggplant seedlings/strain). Non-inoculated plants (mock) served as controls. Percentages in different colours represent different levels of disease severity**. C.** Time-course analysis of disease severity. **D.** Mean relative AUDPC scores (left), average plant fresh weight (center), and fungal re-isolation ratios (right) at the end of the infection experiment (31 d.p.i.). In **C–D**, error bars represent the SE of replicates, and statistical significance of differences between strains was tested by one-way ANOVA followed by Tukey’s post hoc tests. Bars marked with the same letter do not differ significantly (*p*≤0.05).

Our results show that Δ*gtr1* and Δ*tsc2* mutants are defective in regulation of TOR activity and autophagy, which has been previously shown to be important for virulence in plant-pathogenic fungi (Kershaw and Talbot, 2009; Liu et al., 2007). To test whether dysregulation of autophagy could account for the decreased virulence of Δ*gtr1* and Δ*tsc2*, we investigated the pathogenic potential of Δ*atg1*. Our experiments demonstrated that autophagy is indeed essential for full virulence, as Δ*atg1*exhibited strongly reduced disease severity (by 68.1%) and mean AUDPC score (reduced by 85.1%). Moreover, the re-isolation ratio of Δ*atg1* was reduced by 92.8%, indicating a severe defect of this mutant in systemic colonization of the host plant (Supplementary Fig. 4).

## 4. Discussion

The ability of fungi to sense and grow towards environmental resources is instrumental in their growth, development, and survival. In this study we set out to elucidate the molecular mechanisms that govern chemotropic responses during hyphal growth and host interactions of the important plant pathogen *V. dahliae*. To this end, a chemotropism assay on agar plates (Turrà et al., 2015) was optimized for *V. dahliae* enabling reproducible quantification of chemotropic hyphal growth towards a variety of environmental signals.

Our experiments revealed that *V. dahliae* germlings can readily sense and re-orient growth of their hyphal tips towards different carbon and nitrogen sources, similar to what has been previously reported in *F. oxysporum* and *F. graminearum* (Sridhar et al., 2020; Turrà et al., 2015). Nitrogen availability is particularly important for plant pathogenicity (Deng et al., 2015), and the ability of the pathogen to sense and reach suitable nitrogen sources is likely to be important for successful host invasion (Thalineau et al., 2016). Here we found that *V. dahliae* exhibits robust chemotropic responses towards extracellular signals from plant roots. The disease cycle of this fungus normally starts when germination of its resting structures (i.e., microsclerotia) in the soil is triggered by the presence of plant roots nearby, followed by directed hyphal growth towards the roots and invasion. The underlying sensing and chemotropic machinery is therefore essential for pathogenesis. In *F. oxysporum*, which also invades plant roots, chemotropism towards roots is mediated by secreted root peroxidases (Turrà et al., 2015; Nordzieke et al., 2019), and this could be a conserved mechanism for root-invading fungi. Interestingly, our assay also revealed strong chemotropic responses of *V. dahliae* towards the peptide α-pheromone of *F. oxysporum*. Although both fungi are regarded as asexual, our recent investigations showed that *V. dahliae* frequently engages in vegetative hyphal fusions that lead to the formation of viable heterokaryons (Vangalis et al., 2021b). Moreover, genomic evidence for horizontal gene transfer from *F. oxysporum* to *V. dahliae* has been reported (Chen et al., 2018). We speculate that under particular selective conditions the two species might be able to undergo vegetative fusion or communicate via physical cell contact (Haj Hammadeh et al., 2022), and that their pheromone-sensing systems might mediate such interspecific interactions.

The TOR kinase is a central molecular hub that coordinates responses to the availability of various environmental resources in diverse eukaryotes (González and Hall, 2017). In this study, functional characterization of the *V. dahliae* homologs of Gtr1 and Tsc2, two conserved components of the TOR pathway, revealed a role of TOR signaling in sensing and chemotropic growth towards various environmental stimuli. Particularly Gtr1 is important for chemotropic responses towards all the tested stimuli, indicating that the Rag GTPase Gtr1/2 complex plays a central role in the chemosensing process. The mammalian homolog RagA/B-RagC/D is responsible for activation of mTORC1 upon nutrient availability by translocating mTORC1 to the lysosomal surface (Wolfson and Sabatini, 2017). Our results indicate that this function could be conserved in filamentous fungi. On the other hand, Tsc1/2 complex is a TOR repressor that acts under conditions of amino acid deprivation and cellular stress (Gao et al., 2002). Deletion of *V. dahliae tsc2* rendered the fungus unable to re-direct growth towards host-derived signals and the amino acid methionine, but not towards other nutrient sources. Since intracellular amino acid sensors are not conserved across eukaryotes (Wolfson and Sabatini, 2017), filamentous fungi may have evolved different mechanisms, some of which may involve the Tsc1/2 complex. For example, deletion of *tsc1* or *tsc2* in *S. pombe* compromises uptake of arginine and other amino acids, and deletion mutants exhibit decreased expression of amino acid permeases as well as deficiencies in the arginine biosynthesis pathway (Matsumoto et al., 2002; Slegtenhorst et al., 2004). Notably, the Tsc1/2 complex has been implicated in polarization of macrophages upon IL-4 induction (Byles et al., 2013). Together with our results, this points towards a conserved role of the Tsc1/2-TOR system in sensing of a variety of environmental signals and in mediating targeted cellular responses.

In addition to their role in chemotropism, *gtr1* and *tsc2* are required for normal morphogenesis, physiology, and stress response, indicating a conserved role of the TOR signaling pathway in fungal development (Yu et al., 2014). The conidial germination defect of the Δ*tsc2* mutant could be attributed to its inability to undergo autophagy, a process that was shown to be important for conidial germination (Kikuma et al., 2006; Vangalis et al., 2021b). Besides the contributions of Gtr1 and Tsc2 to fungal chemotropism towards root signals and other developmental processes, both components were found to be required for full virulence of *V. dahliae* on eggplant, further supporting a conserved role of TOR signaling in virulence of fungal plant pathogens (Li et al., 2019; Yu et al., 2014). Ιn the case of Tsc2, we hypothesize that its known role in induction of autophagy (Ng et al., 2011), which is further supported by our results, may explain the compromised pathogenicity of the deletion mutant. Consistent with this idea, deletion of *V. dahliae atg1* also resulted in severely decreased virulence, in agreement with findings in other plant pathogens such as *M. oryzae* (Kershaw and Talbot, 2009; Liu et al., 2007). We thus propose that *V. dahliae* Tsc1/2 regulates multiple processes associated with plant pathogenicity, including chemotropism towards roots, fungal development within the host, and autophagy.

To gain a more complete understanding of the signaling components that control *V. dahliae* chemotropism, we used our optimized assay to investigate the role of MAPK signaling. We found that the MAPK Fus3 controls chemotropism towards nutrients, while the GPCR Ste2 and the MAPK Slt2 mediate chemotropic sensing of root signals and α-pheromone. These results are in agreement with previous findings in *F. oxysporum* (Turrà et al., 2015) and reveal that, in contrast to the well-characterized functional association of Ste2 and Fus3 in the yeast mating pathway (Merlini et al., 2013), these asexual fungi employ different associations in processes other than mating, such as Ste2-Slt2 for host and inter-species sensing and Fus3 with unknown receptors for nutrient sensing. In further support of this notion, Fus3 is also functionally uncoupled from Ste2 in conidial and hyphal fusion of filamentous fungi, including *V. dahliae* (Fischer and Glass, 2019; Vangalis et al., 2021a).

In the light of the involvement of both TOR and MAPK signaling in *V. dahliae* chemotropism, an interesting question is whether and how these two pathways interact with each other to control the process. Crosstalk between TOR and MAPK pathways has been reported in fission yeast, where it was proposed to mediate cellular adaptation to multiple environmental cues (Madrid et al., 2016). Ιn *F. oxysporum,* indirect indications of a crosstalk between TOR and the Fus3 homolog, Fmk1, has been reported during regulation of virulence in response to nitrogen sources (López-Berges et al., 2010). Our preliminary results showing altered patterns of Slt2 phosphorylation in the Δ*gtr1* and Δ*tsc2* mutants also point towards a possible crosstalk between TOR and MAPK pathways in chemotropic signaling. We hypothesize that the TOR pathway might be transducing environmental signals to the MAPK cascades for downstream signal processing and cellular responses.

Deletion of *slt2* resulted in a complete absence of microsclerotia and other morphological and physiological defects, as well as increased sensitivity to cell wall stress. These results are consistent with the known role of Slt2 in preserving cell wall integrity, and are in agreement with similar phenotypes reported in other plant pathogens (Jiang et al., 2018). Despite these defects and the role of Slt2 role in sensing of plant roots, deletion of *slt2* had no significant effect on virulence in our assay conditions, and the same result was obtained for the Δ*ste2* mutant. On the contrary, deletion of any of these two components in *F. oxysporum* resulted in reduced virulence (Segorbe et al., 2017; Turrà et al., 2015). Future investigations are required to clarify whether this discrepancy may reflect genetic or regulatory differences between these two species, or whether it may be due to differences in the bioassays used to determine virulence, for example the amount of fungal inoculum. In contrast to Slt2 and Ste2, our results with the Δ*fus3* mutant fully corroborate previous results showing that this MAPK is essential for pathogenicity of *V. dahliae* (Rauyaree et al., 2005), a role that is conserved in all other fungal phytopathogens analyzed so far (Jiang et al., 2018; Turrà et al., 2014).

## 5. Conclusion

Our study establishes the important plant-pathogenic fungus *V. dahliae* as a suitable model for studying the mechanisms of environmental sensing and chemotropism during hyphal growth and host-pathogen interactions. The TOR signaling pathway was shown for the first time to play an important role in chemotropism, along with an involvement in fungal development and virulence. In addition, members of MAPK cascades were found to have conserved roles in chemotropism towards different types of environmental signals, as well as important functions in fungal biology and pathogenicity. Our study contributes to the understanding of the complex chemosensing mechanisms in fungi and raises new questions for future research.

## Acknowledgements

We wish to thank Malena P. Pantou for the Δ*fus3* mutant, which was constructed during her previous work in the laboratory of Milton A. Typas, as well as Melani Mariscal Gómez and Rafael Palos Fernández from the University of Córdoba for their helpful instructions with the plate chemotropism assay.

## Authors contribution statement

**Vasileios Vangalis**: Conceptualization, Methodology, Investigation, Data curation, Formal analysis, Visualization, Writing – original draft, Writing – review and editing.

**Emmanouil A. Markakis**: Methodology, Investigation, Formal analysis, Writing – review and editing.

**Michael Knop**: Funding acquisition, Writing – review and editing.

**Antonio Di Pietro**: Methodology, Supervision, Resources, Funding acquisition, Writing – review and editing.

**Milton A. Typas**: Methodology, Supervision, Resources, Funding acquisition, Writing – review and editing.

**Ioannis A. Papaioannou**: Conceptualization, Methodology, Supervision, Writing – original draft, Writing – review and editing.

## Appendix A. Supplementary Material

Supplementary material related to this article can be found, in the online version, at doi:XXX

## Supplementary Information

**Supplementary Table 1.** *Verticillium dahliae* strains used in this study.

| strain | background | genotype | constructed using plasmid(s) |
| --- | --- | --- | --- |
| 123V | - | wild type | – |
| 123V-Δfus3 | 123V | *fus3::hph* | pUCfus3 |
| 123V-fus3-c | 123V-Δfus3 | *fus3::hph, fus3, neo<sup>R</sup>* | pUCfus3, fus3, pSD1 |
| 123V-Δslt2 | 123V | *Δslt2::neo<sup>R</sup>* | pOSCAR-slt2 |
| 123V-slt2-c | 123V-Δslt2 | *Δslt2::neo<sup>R</sup>, slt2, hph* | pOSCAR-slt2, slt2, pUCATPH |
| 123V-Δfus3 Δslt2 | 123V-Δfus3 | *fus3::hph, Δslt2::neo<sup>R</sup>* | pUCfus3, pOSCAR-slt2 |
| 123V-Δste2 | 123V | *Δste2::neo<sup>R</sup>* | pOSCAR-ste2 |
| 123V-ste2-c | 123V-Δste2 | *Δste2::neo<sup>R</sup>, ste2, hph* | pOSCAR-ste2, ste2, pUCATPH |
| 123V-Δfus3 Δste2 | 123V-Δfus3 | *fus3::hph, Δste2::neo<sup>R</sup>* | pUCfus3, pOSCAR-ste2 |
| 123V-Δgt1 | 123V | *Δgtr1::neo<sup>R</sup>* | pOSCAR-gtr1 |
| 123V-gtr1-c | 123V-Δgtr1 | *Δgtr1::neo<sup>R</sup>, gtr1, hph* | pOSCAR-gtr1, gtr1, pUCATPH |
| 123V-Δtsc2 | 123V | *Δtsc2::hph* | pOSCAR-tsc2 |
| 123V-tsc2-c | 123V-Δtsc2 | *Δtsc2::hph, tsc2, neo<sup>R</sup>* | pOSCAR-tsc2, tsc2, pSD1 |
| 123V-Δatg1 | 123V | *Δatg1::hph* | pOSCAR-atg1 |
| 123V-atg1-c | 123V-Δatg1 | *Δatg1::hph, atg1, neo<sup>R</sup>* | pOSCAR-atg1, atg1, pSD1 |

**Supplementary Table 2.** List of plasmids used in this study.

| plasmid | backbone | description | source |
| --- | --- | --- | --- |
| pUCATPH | - | *hph* | Lu *et al.*, 1994 |
| pSD1 | pBluescript II | $P_{gpdA}$, $P_{trpC}$, $neo^R$ | Nguyen *et al.*, 2008 |
| pOSCAR | pPZP-RCS2 | *A. tumefaciens* binary vector | Paz *et al.*, 2011 |
| pOSCAR-gtr1 | pOSCAR | 5'H<sub>*gtr1*</sub>-$neo^R$-3'H<sub>*gtr1*</sub><br>(H: 2.0 kb-long homology arms) | This study |
| pOSCAR-tsc2 | pOSCAR | 5'H<sub>*tsc2*</sub>-*hph*-3'H<sub>*tsc2*</sub><br>(H: 2.0 kb-long homology arms) | This study |

**Supplementary Table 3.**
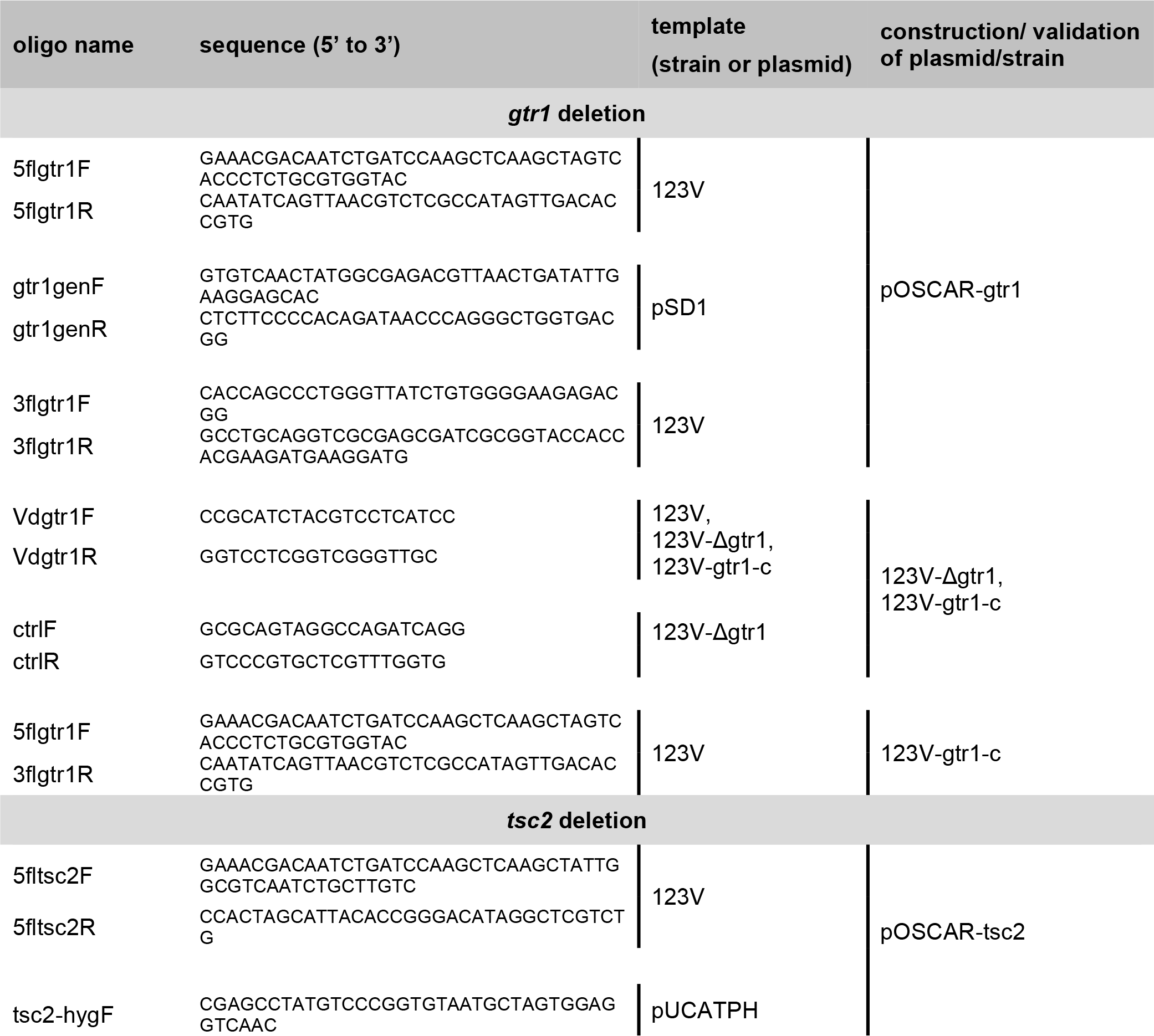
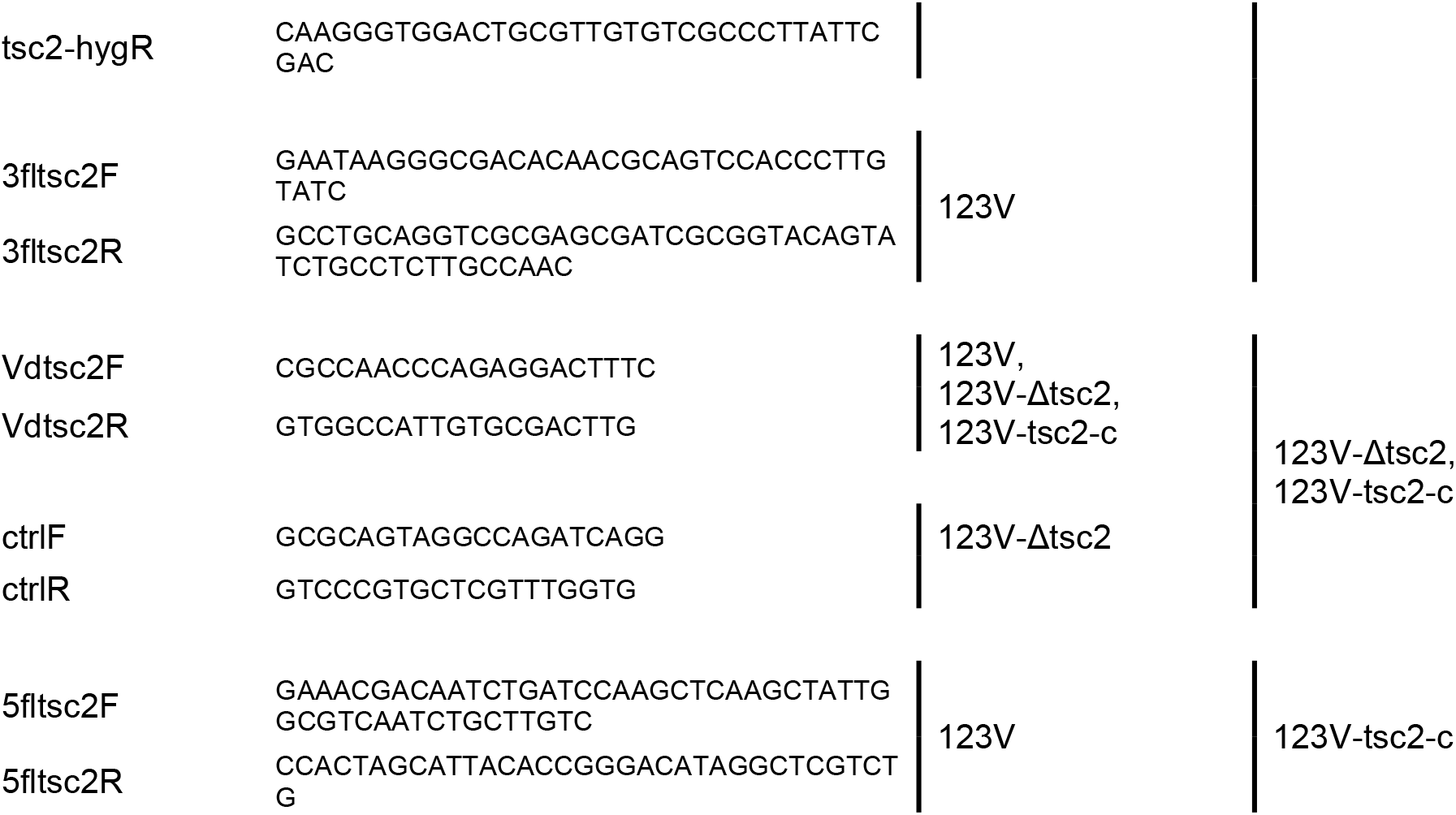
List of DNA oligonucleotides used in this study.

**Supplementary Fig. 1.**
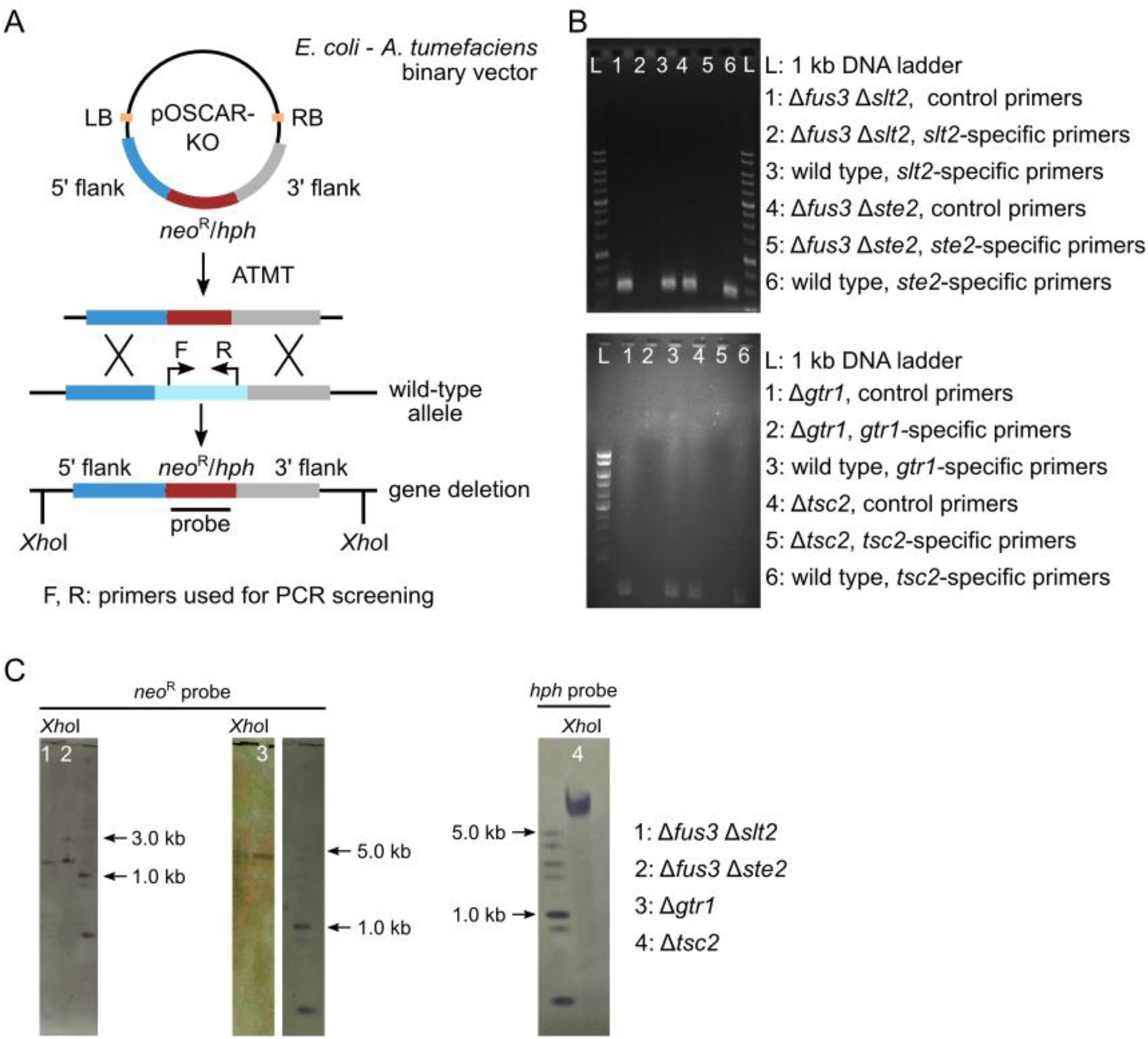
Validation of knockout mutants. **A**. A double homologous recombination-based strategy using *Agrobacterium tumefaciens*-mediated transformation was used for the deletion of the *V. dahliae* 123V homologs of genes *gtr1*, *tsc2*, and the generation of the double knockout mutants Δ*fus3* Δ*slt2* and Δ*fus3* Δ*ste2*. **B**. PCR validation of the deletion mutants Δ*gtr1* and Δ*tsc2* using *gtr1*- or *tsc2*-specific primers as well as the double deletion mutants Δ*fus3* Δ*slt2* and Δ*fus3* Δ*ste2* using *slt2*- or *ste2-*specific primers (Vangalis *et al*., 2021). Primers ctrlF/R (Supplementary Table 3) were used in control reactions for testing template quality. **C**. Validation of mutants by Southern blot after treatment of genomic DNA with *Xho*I, using probes for the *neo*^R^ or the *hph* cassette.

**Supplementary Fig. 2.**
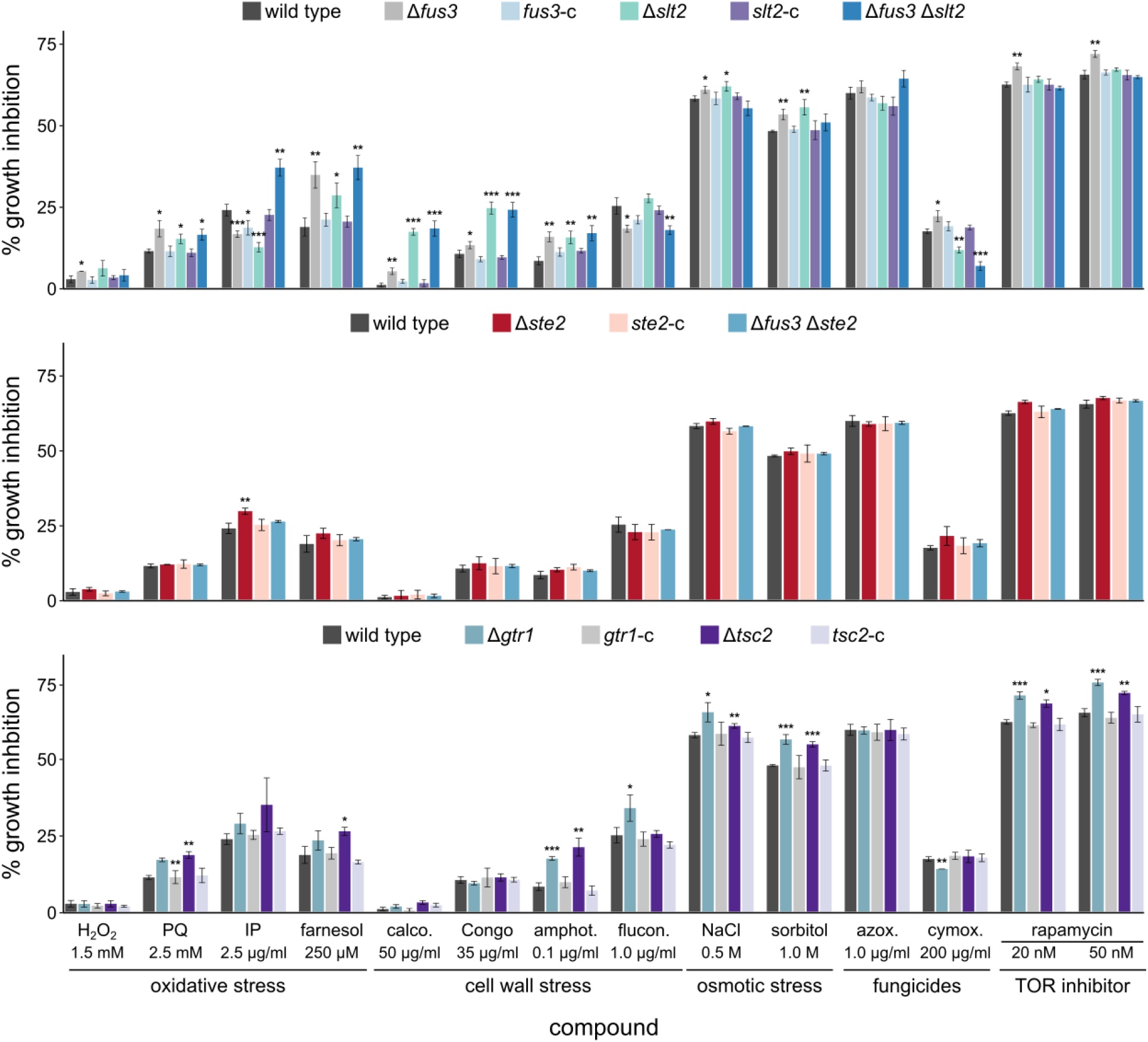
Effect of oxidative and osmotic stresses, cell wall and plasma membrane perturbing agents, and antifungals on the indicated *V. dahliae* strains (**PQ**: paraquat; **azox**: azoxystrobin; **amphot**: amphotericin; **IP**: iprodione; **cymox**: cymoxanil; **calco**: calcofluor white M2R; **Congo**: Congo red; **flucon**: fluconazole). Relative colony growth inhibition was calculated as follows: [(colony diameter on CM − colony diameter in stress condition)/(colony diameter on CM) × 100)]. Bars: SD. Statistical significance of differences to the wild-type strain was tested by Student’s *t*-test (* *p*≤0.05, ** *p*≤0.01, *** *p*≤0.001).

**Supplementary Fig. 3.**
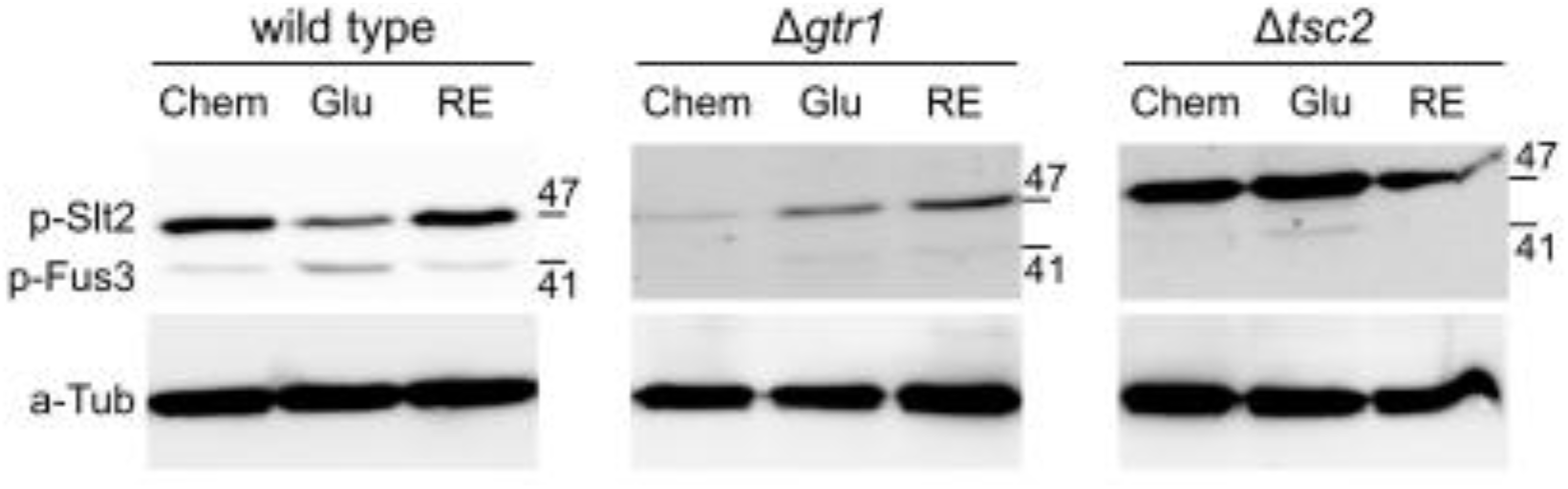
Western blot analysis of Slt2 phosphorylation in the wild-type and the Δ*gtr1* and Δ*tsc2* deletion mutants upon 5 min exposure to glutamate or 10 min exposure to tomato root exudate.

**Supplementary Fig. 4.**
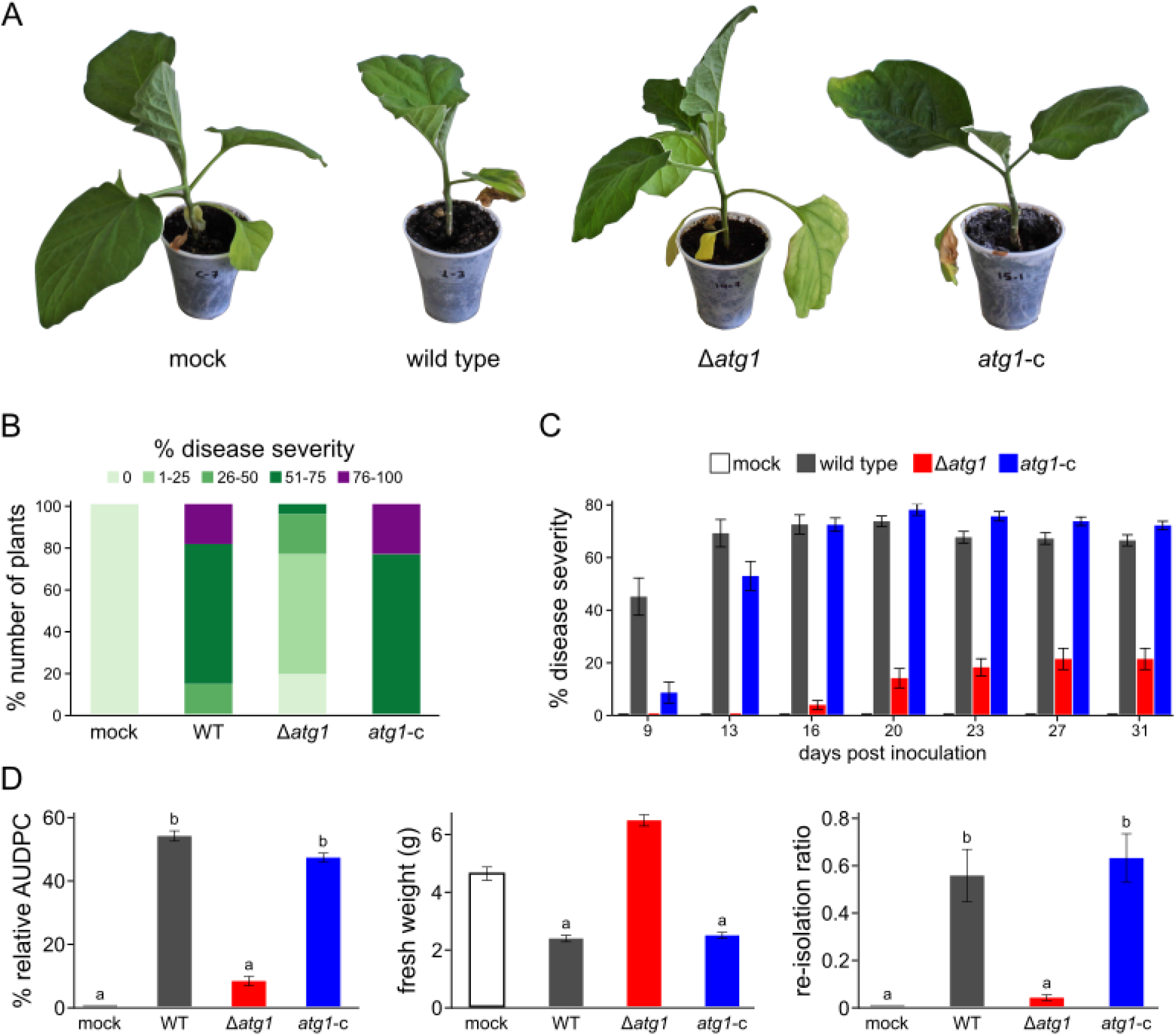
The Δ*atg1* knockout mutant is impaired in pathogenicity. **A**. Representative examples of eggplant plants inoculated with the wild-type strain 123V (WT), the Δ*atg1* deletion mutant and the corresponding complemented strain (*atg1*-c) at 31 days post-inoculation (d.p.i.). **B**. Average disease severity caused by the indicated strains at 31 d.p.i. (21 eggplant seedlings/strain). Non-inoculated plants (mock) served as controls. Percentages in different colours represent different levels of disease severity**. C**. Time-course analysis of disease severity over 31 days. **D**. Average relative AUDPC value of each strain (left), average plant fresh weight (center), and fungal re-isolation ratio at the end of the experiment (31 days). Error bars in **C–D**: SE. Statistical significance of differences between strains was tested by one-way ANOVA followed by Tukey’s post hoc tests. Bars marked with the same letter do not differ significantly (*p*≤0.05).

